# Polycistronic baculovirus expression of SUGT1 enables high-yield production of recombinant leucine-rich repeat proteins and protein complexes

**DOI:** 10.1101/2022.01.19.476911

**Authors:** Kelly Snead, Vanessa Wall, Hannah Ambrose, Dominic Esposito, Matthew Drew

## Abstract

The SHOC2-MRAS-PPP1CA (SMP) complex is a holoenzyme that plays a vital role in the MAP kinase signaling pathway. Previous attempts to produce this challenging three-protein complex have relied on co-infection with multiple viruses and the use of affinity tags to attempt to isolate functional recombinant protein complexes. Leucine-rich repeat containing proteins have been historically challenging to express, and we hypothesized that co-expression of appropriate chaperones may be necessary for optimal production. We describe here how the SUGT1 chaperone can, in conjunction with polycistronic protein expression in baculovirus-infected insect cells, dramatically enhance production yield and quality of recombinant SHOC2, the SMP complex, and other leucine-rich repeat proteins.

**Highlights:**

- Improved yields and purity of LRR proteins and multiprotein complexes through chaperone co-expression, DNA construct design, and use of polycistronic baculovirus.
- Over 300-fold yield increase of SMP protein complex from 0.1 mg/L to 32.6 mg/L.
- SUGT1 co-expression is a highly effective technique for recombinant LRR protein expression and purification.
- Polycistronic baculovirus infection is ideal for production of multiprotein complexes

## 1. Introduction

The developmental diseases Noonan syndrome and Neurofibromatosis type 1 are often caused by germline mutations in the genes encoding RAS and RAS effector proteins, and mutations in these genes are also observed in more than 30% of all human cancers (1). Specific RAS mutations are more prevalent in different types of cancer and there are both upstream and downstream effector proteins that contribute to the development of cancer that are attractive targets for research (2). RAS activates a signaling cascade by binding and activating members of the RAF kinase family, including BRAF and RAF1 proteins. RAF kinases are initially produced in an autoinhibited state in which the adapter protein 14-3-3 is bound in a phosphorylation-dependent manner to two phosphorylated serine residues, which in the case of RAF are pS259 and pS621 (3). In order for signaling through RAS to occur, dephosphorylation of pS259 is required to allow activation of the kinase. A three-protein complex comprised of SHOC2, MRAS, and PPP1CA (known collectively as SMP) has been identified that specifically dephosphorylates RAF1 pS259 and the analogous BRAF pS365 (4). MRAS is a member of the RAS family of proteins and shares regulatory and effector proteins (5). SHOC2 is an evolutionarily well-conserved scaffolding protein comprised primarily of leucine rich repeats (LRRs) (6). Scaffold proteins are important regulators in many signaling pathways and act by binding key proteins into complexes, localizing pathway proteins together, and controlling positive and negative feedback. SHOC2 contributes to modulation of RAF signaling activity through interaction with multiple proteins including RAS, PPP1CA, SCRIB, ERBIN, and HUWE1 (6). SCRIB and ERBIN are LAP (<u>L</u>RR <u>a</u>nd <u>P</u>DZ domain) proteins which mediate SMP activity by interacting with PPP1CA and SHOC2 to disrupt SMP/RAF1 signaling while HUWE1 is involved in ubiquitination of SHOC2 and RAF1 (6,7). The final protein in the complex, PPP1CA, is one of a limited number of serine/threonine phosphatases which can form hundreds of distinct protein-protein complexes to specifically dephosphorylate proteins (8,9). It is unclear how MRAS and SHOC2 target the phosphatase to specifically activate RAF kinases; elucidating the mechanism of this process is important both for understanding a vital RAS signaling component and potentially enabling drug discovery to target this process. Producing large amounts of this complex has been a significant challenge due to poor solubility caused by misfolding of multiple proteins in the complex.

Highly pure, well folded, and functional protein is the goal of recombinant protein expression. Some proteins are highly expressed in soluble form and are easy to purify, while others can be either insoluble or purify as soluble aggregates. Often, expression conditions such as temperature, time of expression, or cell line can enable better protein expression without resorting to altering the target protein. Even after protein expression, the lysis methodology and purification process can make a difference in the yield and quality of the final protein product. However, some proteins cannot be rescued by changing either expression system or purification changes. These proteins need assistance in folding during expression which can only be achieved by co-expressing chaperones and/or co-chaperones. The Heat shock protein 90 (Hsp90) chaperone system is one of the most highly conserved systems among many species and has been extensively studied (10,11). In addition, this system has many co-chaperones that interact with both specific client proteins and the heat shock proteins to promote protein folding and stability. CDC37 is a co-chaperone well known to interact with Hsp90 client kinases, while other co-chaperones such as Hop, p23, and Aha1 act by either stimulating or inhibiting Hsp90 ATPase activity. A study using high throughput LUMIER assays and mass spectrometry identified interaction networks including chaperones, co-chaperones, and client proteins (12). In addition to describing the previously known specificity of the CDC37:kinase interaction, this study found two other domain specific co-chaperones families, Sgt1 and NUDC proteins. Sgt1 is an essential yeast protein (the human homolog is known as SUGT1) and has been found to strongly interact with proteins with LRRs and has a high LUMIER score for interaction with SHOC2 (12,13). Lacking direct interaction with Hsp90, SUGT1 has been implicated in multiple studies to be a co-chaperone for substrate transfer between Hsp70 and Hsp90 in the same way that HOP does for glucocorticoid receptors (13,14). Although the link between SUGT1 and LRR-containing proteins has been known for many years, specific client proteins for SUGT1 have not been extensively reported, especially in the context of recombinant protein expression. The work presented here shows the improvement of SMP complex production using the co-chaperone SUGT1. Co-expression of SUGT1 with SHOC2, PPP1CA, and MRAS resulted in the production of highly pure, functional complexes by both folding SHOC2 and stabilizing PPP1CA to facilitate binding to MRAS. Additionally, we demonstrate the significant benefit of polycistronic co-expression of the SUGT1 chaperone as compared with the more common method of co-infection. We also show that it is possible to isolate similarly pure and high-yield SHOC2:PPP1CA complexes with KRAS4b, HRAS, and NRAS using co-expression of SUGT1. Further testing of SUGT1 co-expression with other LRR proteins such as SCRIB and ERBIN shows significant differences in the way in which SUGT1 might impact protein folding during expression. Overall, these data add another weapon to the armory for high-yield production of challenging protein targets and protein complexes containing the ubiquitous LRR motifs.

## 2. Materials and Methods

### 2.1 DNA

DNA constructs for protein expression were generated using Gateway recombinational cloning or combinatorial Gateway Multisite cloning from a variety of sequence validated Gateway Entry clones (15). In all cases, Entry clones encoding human proteins were produced from synthetic templates constructed by ATUM (Newark, CA) and DNA was optimized for protein expression in insect cells. Templates were flanked by Gateway recombination sites as noted in **Table 1** and fully sequence validated. Several constructs contain protease sites (ENLYFQ/G) for tobacco etch virus (TEV) protease where noted (tev). Final baculovirus expression clones shown in **Table 2** were generated by LR recombination or multisite LR recombination as per the manufacturer’s instructions (Thermo Fisher Scientific, Waltham, MA). Destination vectors include pDest-8 (native baculovirus expression, Thermo Fisher Scientific), pDest-636 (His6-MBP fusion for baculovirus expression, Addgene #159574), and pDest-623 (multisite att4-att3 baculovirus expression, Addgene #161878). After recombination reactions were completed, final expression clones were validated by agarose gel electrophoresis and restriction digest and were used to generate bacmid DNA with the Bac-to-Bac system (Thermo Fisher Scientific, Waltham, MA) using the manufacturer’s instructions.

**Table 1:** Entry clone constructs utilized in this manuscript

| Entry ID | Entry Clone Protein | Gateway sites |
| --- | --- | --- |
| R725-E99 | MRAS(1-178) Q71L | att4-att1r |
| R745-E05 | His6-MBP-tev-PPP1CA(2-330) | att1-att2 |
| R725-E100 | SHOC2 | att2r-att3 |
| R751-E20 | SUGT1 | att4-att5 |
| R745-E30 | MRAS(1-178) Q71L | att5r-att1r |
| R939-E30 | HRAS(1-169) Q61L | att5r-att1r |
| R939-E34 | NRAS(1-169) Q61L | att5r-att1r |
| R949-E92 | KRAS4b(1-169) Q61L | att5r-att1r |
| R745-E50 | tev-SCRIB(1-418) | att1-att2 |
| R745-E51 | tev-ERBIN(1-430) | att1-att2 |
| R751-E14 | SUGT1 | att1-att2 |

**Table 2:** Expression clone constructs utilized in this manuscript

| Clone ID | Protein Expressed | Entry 1 | Entry 2 | Entry 3 | Entry 4 | DV |
| --- | --- | --- | --- | --- | --- | --- |
| R735-M96-623 | SMP complex | R725-E99 | R745-E05 | R725-E100 | N/A | 623 |
| R735-M99-623 | SMP complex + SUGT1 | R751-E20 | R745-E30 | R745-E05 | R725-E100 | 623 |
| R745-M53-623 | SHP complex + SUGT1 | R751-E20 | R939-E30 | R745-E05 | R725-E100 | 623 |
| R745-M57-623 | SNP complex + SUGT1 | R751-E20 | R939-E34 | R745-E05 | R725-E100 | 623 |
| R745-M45-623 | SKP complex + SUGT1 | R751-E20 | R949-E92 | R745-E05 | R725-E100 | 623 |
| R745-X50-636 | His6-MBP-tev-SCRIB | R745-E50 | N/A | N/A | N/A | 636 |
| R745-X51-636 | Hi6-MBP-tev-ERBIN | R745-E51 | N/A | N/A | N/A | 636 |
| R751-X14-008 | SUGT1 | R751-E14 | N/A | N/A | N/A | 008 |

### 2.2 Protein expression in insect cell culture

Baculovirus was produced by transfection of bacmid DNA into Sf9 cells adapted to serum-free suspension and grown in SF900-III SFM media (Thermo Fisher Scientific, Waltham MA). Briefly, 100ml of Sf9 cells were set at 1.5 × 10^6^ cells/ml and transfected using DNA:Cellfectin II (Thermo Fisher Scientific, Waltham MA) lipid complex. The DNA:Cellfectin II complex was prepared using 70ul of prepared bacmid DNA and 250ul Cellfectin II, both diluted in 500ul SF900 III media and mixed together. The transfected culture incubated for 120 hours at 27°C in 250ml Optimum Growth Flasks (Thomson Instrument Company Oceanside, CA) shaking at 105 RPM on an Innova 44 shaker with a 2” orbit (New Brunswick Scientific, Edison, NJ). After harvest of the cell culture supernatant at 120 hours post transfection, the baculovirus titer was measured by qPCR using TaqMan Gene Expression Assay for baculovirus protein GP64 and TaqMan Fast Advanced Master Mix (Thermo Fisher Scientific, Waltham MA).

Protein expression was carried out in Tni-FNL cells isolated and adapted at the Frederick National Laboratory for Cancer Research (16). All proteins were expressed in 1-2 liters of SF900 III SFM media (Thermo Fisher Scientific, Waltham MA) set at 7 × 10^5^ cells/ml in 2.8L or 5L Optimum Growth Flasks (Thomson Instrument Company Oceanside, CA) the day before infection so that cells would double to ∼1.5 × 10^6^ cells/ml on the day of infection for large scale purification. For small scale purification 50mls of Tni-FNL cells were set at 1.5 × 10^6^ cells/ml on the day of infection. Single baculovirus expressions were infected at a multiplicity of infection (MOI) of 3 and multi-baculovirus expressions were infected at MOI of 6 (SHOC2 or SMP) and MOI of 3 (Sgt1). Infected cell culture was incubated for 72 hours at 21°C, shaking at 105 RPM on an Innova 44 shaker with a 2” orbit (New Brunswick Scientific, Edison, NJ). Cells were harvested and cell pellets stored at -80°C until purification.

### 2.3 Protein Purification

Small-scale purification was carried out using a modified version of the procedure first described previously (17). Cell pellets were thawed, resuspended and lysed in 5ml lysis buffer, 20mM HEPES pH7.4, 300mM NaCl, 1mM TCEP, per 50ml of culture. Lysis was performed using the LV-1 Microfluidizer (Microfluidics Corp., Westwood, MA) for 2 passes at 7,000psi on ice. Lysates were then clarified by ultracentrifugation at 100,000 x g for 30 minutes at 4°C. Proteins were purified on the Phynexus MEA (Biotage, Uppsala, Sweden) at 4°C using 40µL bed volume (BV) nickel-charged IMAC tips (Biotage, Uppsala, Sweden). The columns were washed in a 96-well deepwell plate with 20BV dH_2_O and then equilibrated in 20BV of 20mM HEPES pH 7.4, 300mM NaCl, 1mM TCEP, 25mM imidazole. The column tip was loaded with 100µL of sample and then washed with 40BV equilibration buffer. Next, the protein was eluted using 3 × 2BV bump elutions using lysis buffer with 125mM, 250mM, and 500mM imidazole. Total protein, soluble protein, column flowthrough, and column elutions were analyzed by SDS-PAGE and Coomassie-staining.

For large-scale purification, cell pellets were thawed, resuspended and lysed in 100ml lysis buffer, 20mM HEPES pH7.4, 300mM NaCl, 1mM TCEP, per liter of culture. Lysis was performed using the M-110EH Microfluidizer (Microfluidics Corp., Westwood, MA) for 2 passes at 7,000psi on ice. Lysates were then clarified by ultracentrifugation at 100,000 x g for 30 minutes at 4°C and filtered through a 250ml Autofil 0.45µm High Flow PES Bottle Top Filter (Thomas Scientific, Swedesboro NJ) and used immediately. All proteins were purified on NGC chromatography systems (Bio-Rad Laboratories, Hercules CA) using immobilized metal affinity chromatography (IMAC) and size exclusion chromatography (SEC). The flowrate for all IMAC steps was 5ml/min and for SEC was 1ml/min. Clarified lysate was adjusted to 35mM imidazole and loaded onto a 5ml Ni Sepharose High Performance nickel-charged column (GE Healthcare, Chicago, IL) equilibrated in 20mM HEPES pH 7.4, 300mM NaCl, 1mM TCEP, 35mM imidazole. The column was washed for 5 column volumes (CV) with the equilibration buffer and protein was eluted using a gradient of 7-100% Buffer B (20mM HEPES pH 7.4, 300mM NaCl, 1mM TCEP, 500mM imidazole) for 20 CV, and column elution collected in 5 ml fractions. Total protein, soluble protein, and elution fractions were analyzed by SDS-PAGE and Coomassie-staining. Appropriate fractions were pooled and dialyzed into 2 liters 20mM HEPES pH 7.4, 300mM NaCl, 1mM TCEP with His6-TEV protease at a 1:20 (v/v) protease:pool volume ratio overnight at 4°C. The digested material was processed by a second IMAC to separate the cleaved His6-MBP tag on the PPP1CA from the SMP complex. The second IMAC column was equilibrated with 20mM HEPES pH7.4, 300mM NaCl, 1mM TCEP and proteins were eluted with a shallow gradient from 0-10% Buffer B for 10CV followed by a 10-100% Buffer B gradient for 10CV. Elution fractions containing the desired complex by Coomassie stained SDS-PAGE were pooled from the shallow gradient and concentrated using a 30K Amicon Ultra 15 Centrifugal Filter Units (Millipore Sigma, Burlington MA) down to 5ml for size exclusion chromatography. The concentrated sample was applied to an SEC column (HiLoad 16/600 Superdex 200, Cytiva, Chicago IL) equilibrated in final buffer (20mM HEPES pH 7.4, 150mM NaCl, 1mM TCEP). Final buffer was flowed over the column at 1.2 ml/min and fractions were pooled based on Coomassie-stained SDS-PAGE and a final yield concentration was determined by measuring A_280_ (Nanodrop 2000C spectrophotometer (Thermo Fisher Scientific, Waltham MA).

## 2. Results/Discussion

### 3.1 Optimization of expression constructs for SMP complex production

Initial SHOC2, MRAS, and PPP1CA constructs (**Fig. 1A**) tested were designed with three different purification tags to be used with multiple affinity purifications for high-purity final protein. The glutathione-S-transferase (GST) tagged MRAS had good levels of expression and high solubility. However, both the TwinStrep-tagged SHOC2 and His6-tagged PPP1CA had high levels of expression but were almost completely insoluble in our initial tests (data not shown) leading to production of a complex at very low yield (0.1 mg/L) and poor purity (**Fig. 1C**, left panel)

**Fig. 1.**
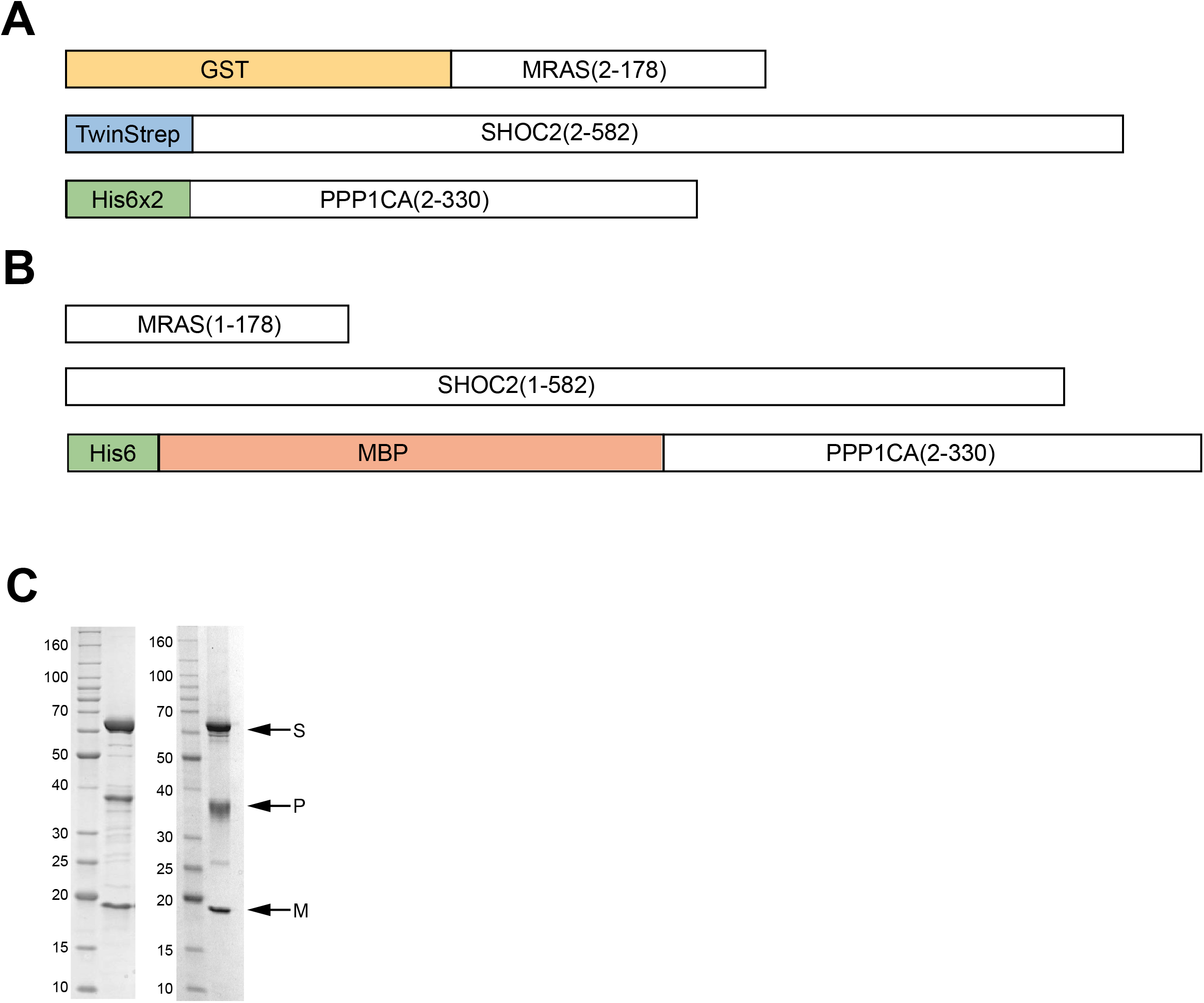
Optimization of constructs for SMP protein complex production. A. Initial constructs tested included a GST tag (yellow) on MRAS, a TwinStrep tag (blue) on SHOC2, and a His6×2 tag (green) on PPP1CA, all containing a tobacco etch virus protease site between the tag and protein of interest. B. Optimized constructs have untagged MRAS and SHOC2 and a single His6 tag (green) with a maltose binding protein (MBP, red) on PPP1CA, separated by a tobacco etch virus protease site. C. SDS-PAGE/Coomassie staining analysis of final purified proteins for initial constructs (left) and optimized constructs (right). Protein bands representing SHOC2 (S), MRAS (M), and PPP1CA (P) are noted with arrows. Molecular weight markers are shown in kilodaltons.

Maltose binding protein (MBP) is a common solubility enhancing tag used in both prokaryotic and eukaryotic expression systems (18). Based on success with other proteins in our laboratory, we redesigned all of the SMP constructs to resemble our standard MBP-based protein expression strategy more closely. Placing MBP tags on all three components improved the apparent solubility of each protein but the overall yield of purified complex was not significantly improved (data not shown). After testing multiple tags and expression conditions using these constructs, we settled on an optimal combination of native full-length SHOC2, native MRAS G-domain (amino acids 1-178, lacking the C-terminal hypervariable tail), and full-length His6-MBP-tev-PPP1CA containing a tobacco etch virus (TEV) protease site (tev) between the fusion tag and PPP1CA. (**Fig. 1B**). Using this combination of constructs, we were able to increase both the yield and purity of the complex (**Table 3, Fig. 1C**) from 0.1 mg/L to 2 mg/L. However, purity levels were still insufficient for downstream structural biology needs, and the triple infection required for production of the complex was suboptimal for final complex yield and reproducibility.

**Table 3:** Purification yields of SHOC2-MRAS-PPP1CA complexes

| Proteins | Viruses used | Protein Yield (mg/L) | Avg. Yield (mg/L) |
| --- | --- | --- | --- |
| SHOC2 + MRAS + PPP1CA | 3 | 3.0, 1.9, 1.1, 2.1 | 2.0 ± 0.7 |
| SHOC2 + MRAS + PPP1CA + SUGT1 | 4 | 5.8, 3.7, 3.7 | 4.4 ± 1.0 |
| SHOC2/MRAS/PPP1CA | 1 | 5.5, 5.7, 3.5, 4.6 | 4.8 ± 0.9 |
| SHOC2/MRAS/PPP1CA + SUGT1 | 2 | 26.5, 27.0, 27.0 | 26.8 ± 0.2 |
| SHOC2/MRAS/PPP1CA/SUGT1 | 1 | 33.1, 37.5, 27.2 | 32.6 ± 4.2 |

### 3.2 SUGT1 co-expression with SHOC2 and SMP complex

Several literature reports suggest that chaperone/co-chaperone systems specific to a variety of protein domains, such as leucine-rich repeats like those found in SHOC2, were vital to the proper intracellular production of correctly folded protein complexes (12,13). We hypothesized that co-expression of co-chaperones might enable higher yield and solubility of our highly overexpressed recombinant proteins. Initially, we tested co-expression of human SUGT1 (a homolog of yeast SGT1) on the production of SHOC2. **Fig. 2A** shows total and soluble protein expression and microscale purification of His6-MBP-SHOC2 alone (left) and in combination with SUGT1 (right). This test clearly showed a dramatic improvement in both SHOC2 solubility and purification yield. Expression of SHOC2 in the absence of SUGT1 produced less than ∼20% soluble protein, while addition of the co-chaperone led to 100% soluble SHOC2. Based on this result we performed a quadruple infection with SUGT1 and all three components of the SMP complex (**Fig. 1B**), which more than doubled the final yield of the complex from 2 mg/L to 4.4 mg/L (**Fig. 2B, Table 3**). Pooled fractions from this purification were assessed by preparative size exclusion chromatography (SEC), and based on known molecular weight standards, the complex migrated as expected for a 122 kDa protein complex (**Fig. 2C**).

**Fig. 2.**
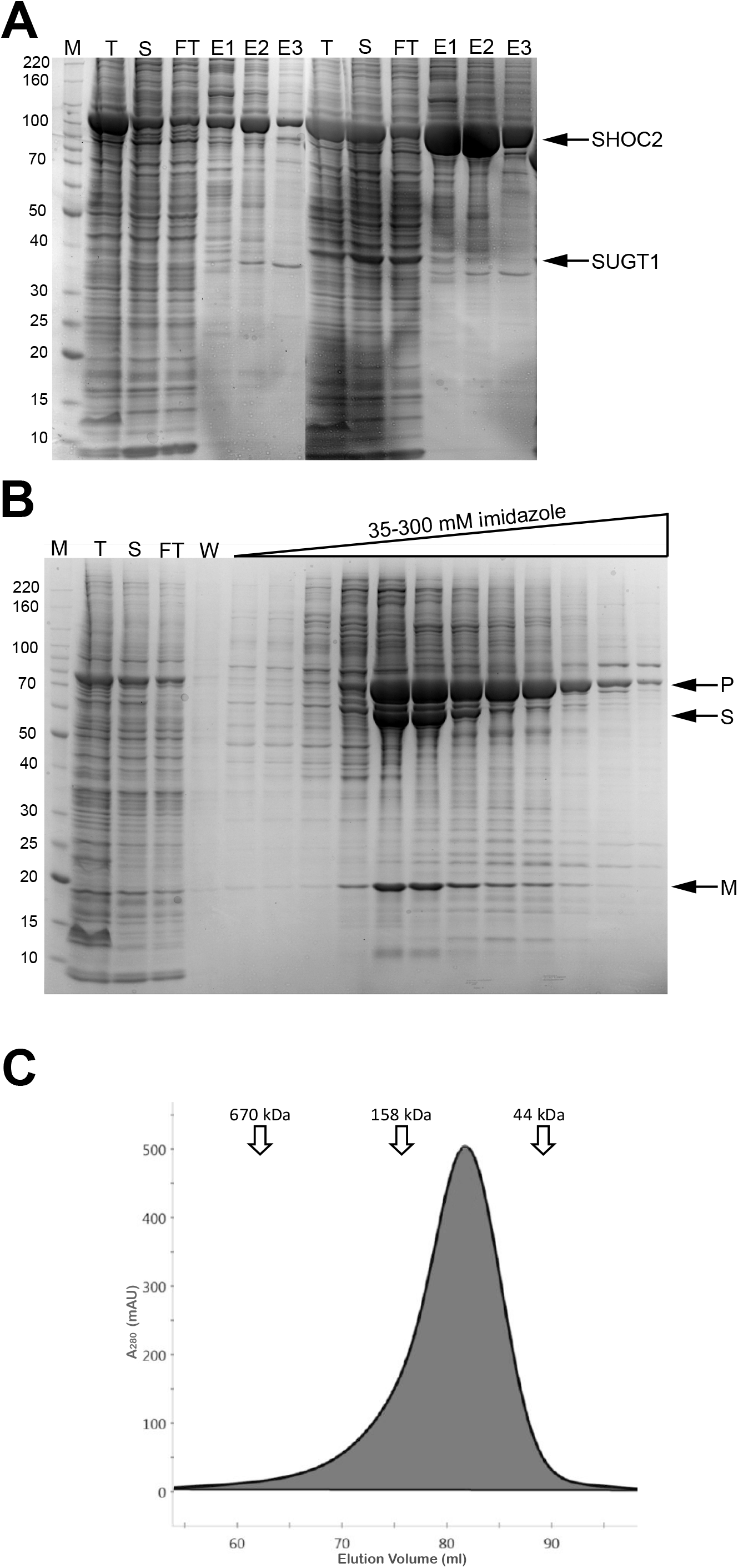
SUGT1 co-expression improves SHOC2 and SMP complex production. A. SDS-PAGE/Coomassie staining analysis of small scale IMAC chromatography of full length SHOC2 alone (left) and with SUGT1 co-expression (right), M – molecular weight standards noted in kilodaltons, T – total protein expression, S – soluble protein/column load, FT – column flowthrough. E1 – 125mM imidazole elution fraction, E2 – 250mM imidazole elution fraction, E3 – 500mM imidazole elution fraction. B. SDS-PAGE/Coomassie staining analysis of large scale IMAC chromatography of SMP and SUGT1 quadruple infection, W – column wash. C. SEC chromatogram of SMP and SUGT1 quadruple infection with molecular weight standards noted at appropriate elution volumes in kilodaltons.

### 3.3 Polycistronic expression of the SMP protein complex

The multiplicity of infection (MOI) for a baculovirus infection is a vital parameter in recombinant protein expression in insect cells, particularly when infecting cells with multiple baculoviruses. The *Trichoplusia ni* Tni-FNL cell line supports high protein expression, but relatively low virus amplification compared to Sf9 cells (16,19). The reduced virus amplification capacity of Tni-FNL cells means that an MOI of at least 3 is required to obtain at least one viral particle per cell for greater than 95% of the cells in culture (20). It is especially crucial when expressing multiple proteins that form a complex, that the correct MOI ratio between each virus is determined to permit expression of each protein in equimolar amounts. However, during a multi-baculovirus infection it is nearly impossible to ensure that each cell is infected with the same amount of each virus or that each cell is infected with at least one copy of each virus. The result is a heterogenous sample of cells which may express 1, 2, or 3 proteins at varying levels, which can lead to significant accumulation of failed complexes or insoluble proteins. To solve this problem, many laboratories have developed methods to generate polycistronic baculoviruses which express multiple genes from one virus (21,22). In our laboratory, we have addressed this problem using multisite Gateway recombinational cloning to generate polycistronic constructs in which each gene is transcribed by its own baculovirus polyhedrin promoter in a single viral genome (15).

Using a single baculovirus with the optimal SHOC2, MRAS, and PPP1CA constructs increased the yield of the SMP complex from 2 mg/L to 4.8 mg/L, more than doubling the yield compared with the triple infection (**Table 3, Fig. 3A**). This yield improvement likely resulted from every cell now expressing all three proteins in equal amounts. Further co-infection of the SMP polycistronic baculovirus with a virus containing SUGT1 increased the yield an additional 6-fold to 26.8 mg/L (**Fig. 3C)**. Even though both the SMP polycistronic virus alone and with SUGT1 produced very similar amounts of protein in the initial cell extracts (“T” lanes, **Fig. 3A** and **Fig. 3C**), SMP complex formation and yield without SUGT1 is highly limited by the reduced levels of properly folded SHOC2 and stable PPP1CA. With SUGT1 co-infection, the majority of the PPP1CA is folded into the complex and the SHOC2 is removed from the flowthrough of the IMAC purification (“FT” lanes, **Fig. 3A** and **Fig. 3C**).

**Fig. 3.**
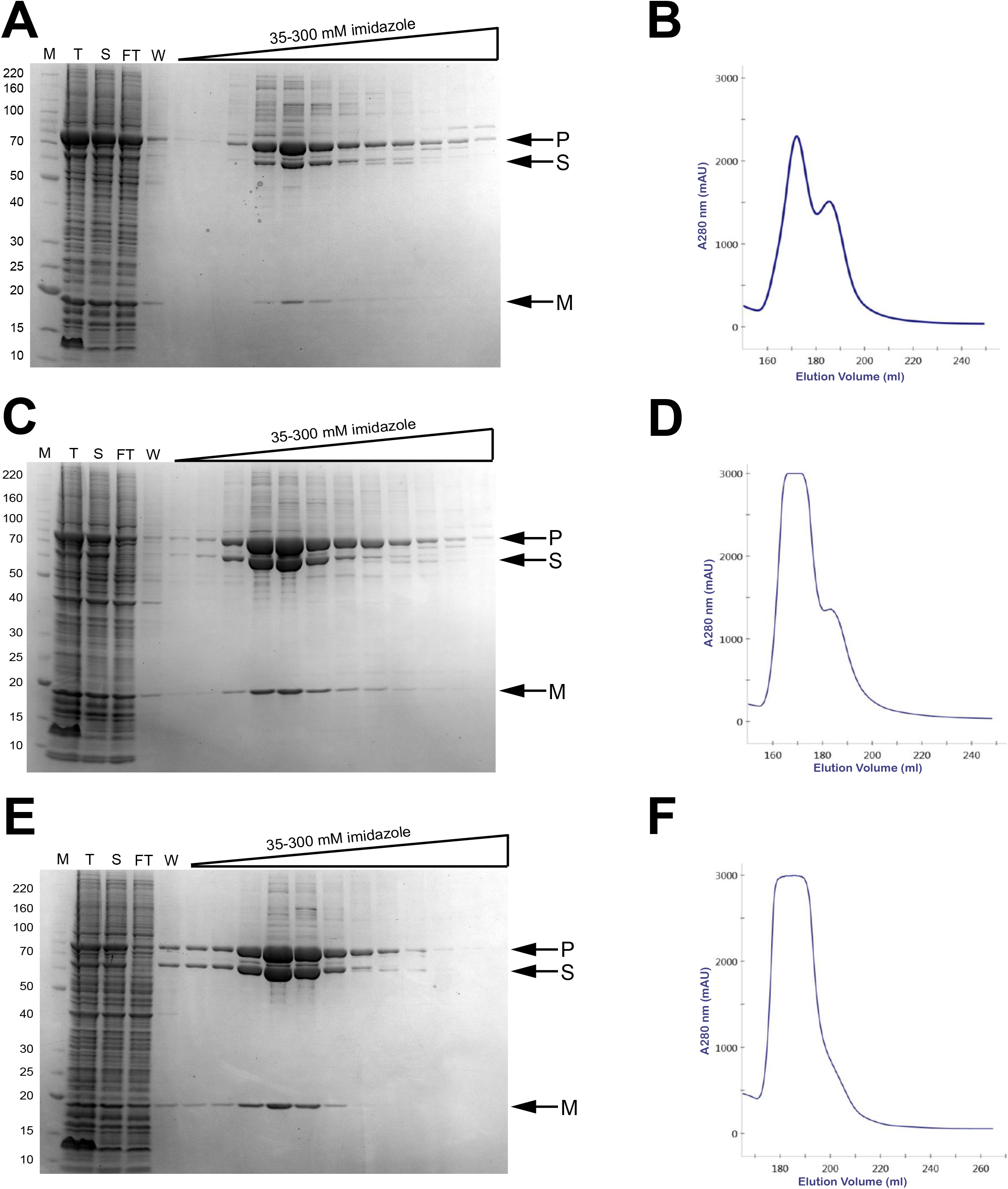
Polycistronic production of SUGT1 and SMP complexes further improves yield and quality of the SMP complex. A. SDS-PAGE/Coomassie staining analysis of large scale IMAC chromatography of polycistronic SMP without SUGT1. B. IMAC chromatogram of polycistronic SMP without SUGT1. C. SDS-PAGE/Coomassie staining analysis of large scale IMAC chromatography of polycistronic SMP with co-infection of SUGT1. D. IMAC chromatogram of polycistronic SMP with co-infection of SUGT1. E. SDS-PAGE/Coomassie staining analysis of large scale IMAC chromatography of polycistronic SMP including SUGT1. F. IMAC chromatogram of polycistronic SMP including SUGT1. M – molecular weight standards noted in kilodaltons, T – total protein expression, S – soluble protein/column load, FT – column flowthrough, W – column wash.

To further improve the system, we combined the three SMP proteins and SUGT1 all on a single polycistronic baculovirus. With this new virus, we were able to generate SMP complex at a yield of 32.6mg/L (**Table 3). Fig. 3E** shows the purification of this complex and notably demonstrates that nearly all SHOC2 and PPP1CA protein is now present in the proper SMP complex. The IMAC chromatograms (**Fig. 3D** as compared with **Fig. 3B**) also highlight how the presence of SUGT1 increases the purifiability of the complex while still leaving some excess PPP1CA uncomplexed in the righthand peak. This is likely due to the unequal ratio of viruses often observed in co-infection, which can lead to incomplete complex formation in some cells. In the polycistronic virus which expresses all 4 proteins from the same virus (**Fig. 3F**), this peak is nearly eliminated, suggesting that most of the purified protein represents the full SMP complex. In total, the combination of vector, expression, and purification optimization increased the overall yield of SMP complex by over 300-fold from 0.1mg/L to 32.6mg/L, and dramatically improved purity of the final complex.

Purified SMP complexes were validated in a dephosphorylation assay which showed that they could remove the specific phosphoserine residues on RAF1 and BRAF without any effect on other phosphoserines (data not shown), indicating that the complexes produced were functional (or active). In addition, the high yield of extremely pure proteins produced using this method led to the generation of crystals of both SHOC2 and SMP complex which are currently being used to elucidate the high-resolution crystal structure of these proteins for the first time (Bonsor D., et al, manuscript in preparation).

### 3.4 SUGT1 co-expression with other SHOC2-containing protein complexes

In addition to the SMP complex, we also produced SHOC2/PPP1CA complexes with other RAS isoforms, including KRAS (SKP), HRAS (SHP), and NRAS (SNP). Initially, in the absence of SUGT1 co-expression, we were unable to produc significant quantities of these complexes, which are believed to have lower affinity than the SMP complex. However, when combined in the 4-piece polycistronic baculovirus format used for SMP, we were able to isolate at high yield the same 1:1:1 functional complex as observed with SMP (**Fig. 4**). As expected from the lower affinity of the complexes, the yields were significantly lower than we saw with the SMP complex (**Table 4**), but still high enough to produce sufficient high-quality protein for biochemical and structural experiments.

**Fig. 4.**
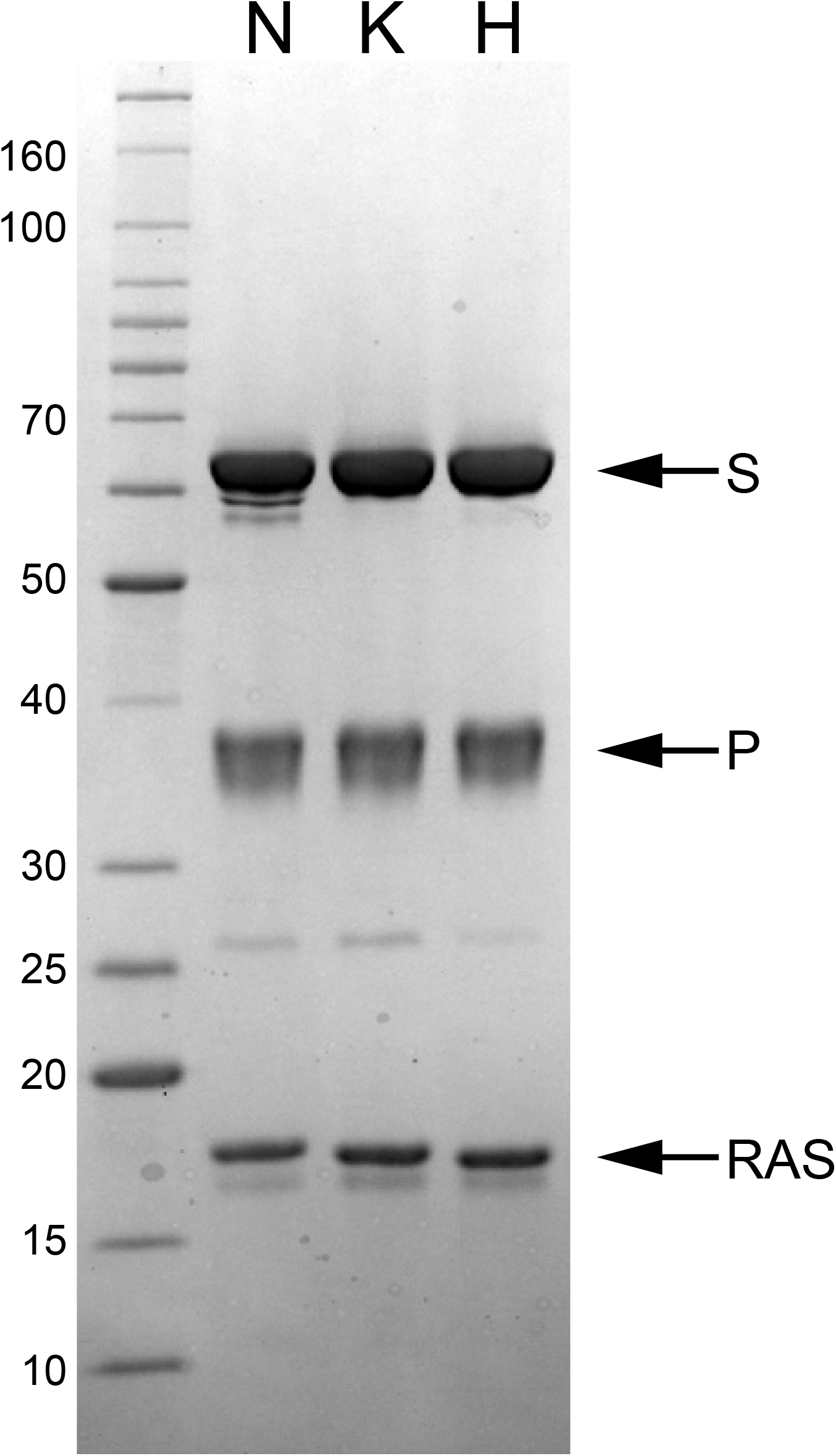
SHOC2/PPP1CA complexes with other RAS isoforms. SDS-PAGE/Coomassie staining analysis of final purified proteins for SNP (left), SKP (middle), and SHP (right) complexes. Each complex was made using the same SHOC2, PPP1CA and SUGT1 polycistronic construct with various RAS G-domain isoforms as noted.

**Table 4:** Purification yields of SHOC2-RAS-PPP1CA complexes with different RAS isoforms

| Construct | RAS isoform | Yield (mg/L) | Average (mg/L) |
| --- | --- | --- | --- |
| SMP | MRAS | 33.1, 37.5, 27.2 | 32.6 ± 4.2 |
| SNP | NRAS | 18.5, 23.5, 22.2 | 21.4 ± 2.1 |
| SKP | KRAS | 11.1, 10.6, 14.5 | 12.1 ± 1.7 |
| SHP | HRAS | 12.3, 23.2, 24.7 | 20.0 ± 5.5 |

### 3.5 SUGT1 co-expression with other LRR-containing proteins

SCRIB is another LRR-containing protein that is poorly folded when expressed recombinantly in insect cells. We co-expressed SCRIB with SUGT1 to determine experimentally whether the co-chaperone might have a similar impact on SCRIB as it did on SHOC2. As shown in **Fig. 5A**, expression of SCRIB in the absence of SUGT1 produces high levels of soluble protein, but the majority of the protein fails to bind the IMAC column. This likely represents soluble aggregate and/or misfolded protein due to the lack of co-chaperone for SCRIB folding. When SUGT1 is co-expressed using a second virus (**Fig. 5B**), binding of SCRIB to the IMAC column is enhanced. Although the protein is highly expressed both with and without SUGT1, the final yield increased from 9.3 mg/L to 26.4 mg/L despite the suboptimal use of co-infection instead of co-expression of SUGT1.

**Fig. 5.**
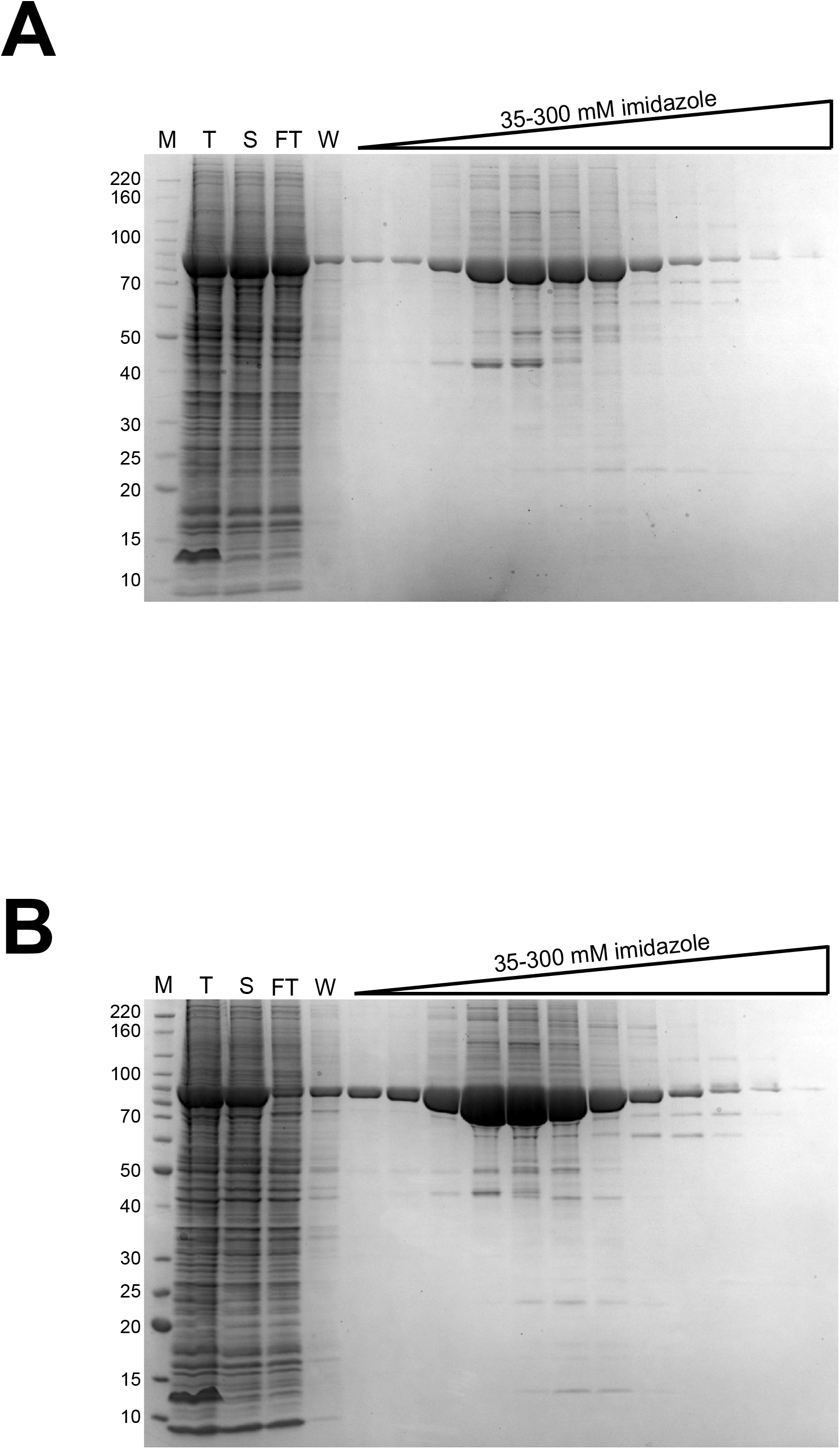
Effect of SUGT1 co-expression on production of SCRIB. The SCRIB construct used for testing co-expression with SUGT1 contains the LRR domains including amino acids 1-418 and excludes other domains. A. SDS-PAGE/Coomassie staining analysis of IMAC of SCRIB (1-418) alone. B. SDS-PAGE/Coomassie staining analysis of IMAC of SCRIB (1-418) with SUGT1 co-expression by co-infection. M – molecular weight standards noted in kilodaltons, T – total protein expression, S – soluble protein/column load, FT – column flowthrough, W – column wash.

ERBIN is an LRR-containing protein involved in the regulation of the activity of the SMP complex. Expressed alone (**Fig. 6A** and **Fig. 6B**), ERBIN is mostly aggregated, and solubility tag cleavage is poor (“Pool 1”), with only a small percentage (“Pool 2”) of monomeric protein that purifies with many higher molecular weight contaminants. With co-expression of SUGT1 (**Fig. 6C** and **Fig. 6D**), there is an additional aggregate pool that co-elutes with SUGT1 and cleaved protein (“Pool 2”), a pool that is cleaved ERBIN with SUGT1 (“Pool 3”), and the monomer pool (“Pool 4”). This collection of different fractions on the SEC suggests that the SUGT1 binds ERBIN but gets stuck and cannot cycle completely through the chaperone system to fold the remainder of the aggregated protein. It is possible that additional co-chaperones could be needed to fully allow ERBIN to fold completely. In comparison to SHOC2, both SCRIB and ERBIN have many more predicted disordered regions and additional functional domains which could account for the difference in the ability of SUGT1 to completely rescue the production of these proteins.

**Fig. 6.**
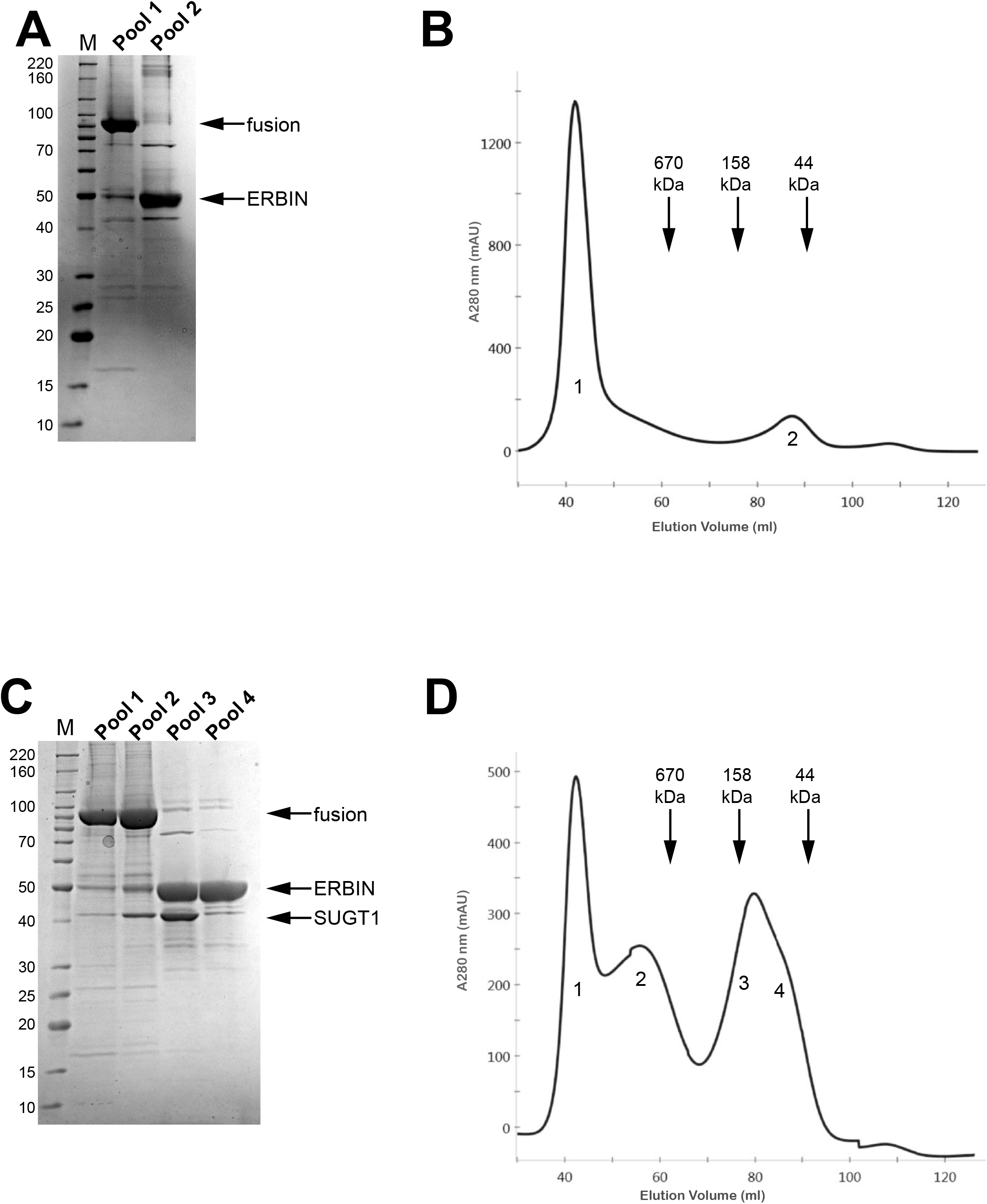
Effect of SUGT1 co-expression on production of ERBIN. The ERBIN construct used for testing co-expression with SUGT1 contains the LRR domains including amino acids 1-430 and excludes other domains. A. SDS-PAGE/Coomassie staining analysis of the SEC aggregate peak (pool 1) and included peak (pool 2) for ERBIN alone. B. SEC chromatogram of ERBIN alone. C. SDS-PAGE/Coomassie staining analysis of the SEC aggregate peaks (pool 1 and pool 2) and included peaks (pool 3 and pool 4) for ERBIN with SUGT1 co-expression. D. SEC chromatogram of ERBIN with SUGT1 co-expression. Molecular weight standards (M) on SDS-PAGE gels are noted in kilodaltons. Locations of molecular weight standard elution on SEC columns are denoted by vertical arrows and relative sizes are indicated in kilodaltons.

## 4. Conclusions

We have demonstrated the ability to significantly improve yields of recombinant protein complexes containing LRR proteins using a combination of construct design, co-chaperone co-expression using SUGT1, and polycistronic baculoviruses. It is likely that for these proteins, any enhancement in production utilizing solubility tags only increases the level of soluble aggregate formation, and in the absence of the proper co-chaperone folding system, production of high-yield and high-purity proteins would be challenging. While the largest improvement seen was in the production of the SMP complex, we also have shown that other proteins in the LRR family such as ERBIN and SCRIB are also improved to different levels using this SUGT1 system. It remains to be seen why some proteins may perform better than others or whether some proteins may require additional chaperones to complete their proper folding. However, it is clear that SUGT1 plays a major role in the folding process, and it provides a significant weapon in the arsenal of chaperone-mediated recombinant protein production which should be considered for the production of many important proteins that contain leucine-rich repeats. In addition, we have provided another example that the optimal production of protein complexes in baculovirus-infected insect cells benefits significantly from polycistronic viruses that ensure proper expression of all proteins in each cell. As structural biology and drug discovery efforts move consistently towards research into larger multiprotein complexes, the need to ensure consistent high-quality production of these proteins will continue to grow.

## Abbreviations

CV: column volume
IMAC: immobilized metal ion affinity chromatography
LRR: leucine-rich repeat
MWCO: molecular weight cut-off
SDS-PAGE: sodium dodecyl sulfate-polyacrylamide gel electrophoresis
SEC: size exclusion chromatography
SMP: SHOC2/MRAS/PPP1CA protein complex
TEV: tobacco etch virus proteins

## Acknowledgements

We acknowledge our colleagues Carissa Grose and Simon Messing from the Protein Expression Laboratory at the Frederick National Laboratory for technical support, and Bill Gillette for critical review of the manuscript. Initial SMP expression constructs were kindly provided by Dr. Pablo Rodriguez-Viciana from University College London. This project has been funded in whole or in part with Federal funds from the National Cancer Institute, National Institutes of Health, under contract number 75N91019D00024. The content of this publication does not necessarily reflect the views or policies of the Department of Health and Human Services, nor does mention of trade names, commercial products, or organizations imply endorsement by the U.S. Government.

## References

1. Schubbert, S., Shannon, K., and Bollag, G. (2007) Hyperactive Ras in developmental disorders and cancer. Nat Rev Cancer 7, 295–308

2. Stephen, A. G., Esposito, D., Bagni, R. K., and McCormick, F. (2014) Dragging ras back in the ring. Cancer Cell 25, 272–281

3. Terrell, E. M., and Morrison, D. K. (2019) Ras-Mediated Activation of the Raf Family Kinases. Cold Spring Harb Perspect Med 9

4. Rodriguez-Viciana, P., Oses-Prieto, J., Burlingame, A., Fried, M., and McCormick, F. (2006) A phosphatase holoenzyme comprised of Shoc2/Sur8 and the catalytic subunit of PP1 functions as an M-Ras effector to modulate Raf activity. Mol Cell 22, 217–230

5. Young, L. C., and Rodriguez-Viciana, P. (2018) MRAS: A Close but Understudied Member of the RAS Family. Cold Spring Harb Perspect Med 8

6. Jang, H., Stevens, P., Gao, T., and Galperin, E. (2021) The leucine-rich repeat signaling scaffolds Shoc2 and Erbin: cellular mechanism and role in disease. FEBS J 288, 721–739

7. Young, L. C., Hartig, N., Munoz-Alegre, M., Oses-Prieto, J. A., Durdu, S., Bender, S., Vijayakumar, V., Vietri Rudan, M., Gewinner, C., Henderson, S., Jathoul, A. P., Ghatrora, R., Lythgoe, M. F., Burlingame, A. L., and Rodriguez-Viciana, P. (2013) An MRAS, SHOC2, and SCRIB complex coordinates ERK pathway activation with polarity and tumorigenic growth. Mol Cell 52, 679–692

8. Bollen, M., Peti, W., Ragusa, M. J., and Beullens, M. (2010) The extended PP1 toolkit: designed to create specificity. Trends Biochem Sci 35, 450–458

9. Korrodi-Gregorio, L., Esteves, S. L., and Fardilha, M. (2014) Protein phosphatase 1 catalytic isoforms: specificity toward interacting proteins. Transl Res 164, 366–391

10. Li, J., Soroka, J., and Buchner, J. (2012) The Hsp90 chaperone machinery: conformational dynamics and regulation by co-chaperones. Biochim Biophys Acta 1823, 624–635

11. Sahasrabudhe, P., Rohrberg, J., Biebl, M. M., Rutz, D. A., and Buchner, J. (2017) The Plasticity of the Hsp90 Co-chaperone System. Mol Cell 67, 947–961 e945

12. Taipale, M., Tucker, G., Peng, J., Krykbaeva, I., Lin, Z. Y., Larsen, B., Choi, H., Berger, B., Gingras, A. C., and Lindquist, S. (2014) A quantitative chaperone interaction network reveals the architecture of cellular protein homeostasis pathways. Cell 158, 434–448

13. Stuttmann, J., Parker, J. E., and Noel, L. D. (2008) Staying in the fold: The SGT1/chaperone machinery in maintenance and evolution of leucine-rich repeat proteins. Plant Signal Behav 3, 283–285

14. Bhattacharya, K., Weidenauer, L., Luengo, T. M., Pieters, E. C., Echeverria, P. C., Bernasconi, L., Wider, D., Sadian, Y., Koopman, M. B., Villemin, M., Bauer, C., Rudiger, S. G. D., Quadroni, M., and Picard, D. (2020) The Hsp70-Hsp90 co-chaperone Hop/Stip1 shifts the proteostatic balance from folding towards degradation. Nat Commun 11, 5975

15. Wall, V. E., Garvey, L. A., Mehalko, J. L., Procter, L. V., and Esposito, D. (2014) Combinatorial assembly of clone libraries using site-specific recombination. Methods Mol Biol 1116, 193–208

16. Talsania, K., Mehta, M., Raley, C., Kriga, Y., Gowda, S., Grose, C., Drew, M., Roberts, V., Cheng, K. T., Burkett, S., Oeser, S., Stephens, R., Soppet, D., Chen, X., Kumar, P., German, O., Smirnova, T., Hautman, C., Shetty, J., Tran, B., Zhao, Y., and Esposito, D. (2019) Genome Assembly and Annotation of the Trichoplusia ni Tni-FNL Insect Cell Line Enabled by Long-Read Technologies. Genes (Basel) 10

17. Gillette, W. K., Esposito, D., Taylor, T. E., Hopkins, R. F., Bagni, R. K., and Hartley, J. L. (2011) Purify First: rapid expression and purification of proteins from XMRV. Protein Expr Purif 76, 238–247

18. Sun, P., Tropea, J. E., and Waugh, D. S. (2011) Enhancing the solubility of recombinant proteins in Escherichia coli by using hexahistidine-tagged maltose-binding protein as a fusion partner. Methods Mol Biol 705, 259–274

19. Wilde, M., Klausberger, M., Palmberger, D., Ernst, W., and Grabherr, R. (2014) Tnao38, high five and Sf9--evaluation of host-virus interactions in three different insect cell lines: baculovirus production and recombinant protein expression. Biotechnol Lett 36, 743–749

20. Palomares, L. A., Mena, J. A., and Ramirez, O. T. (2012) Simultaneous expression of recombinant proteins in the insect cell-baculovirus system: production of virus-like particles. Methods 56, 389–395

21. Pelosse, M., Crocker, H., Gorda, B., Lemaire, P., Rauch, J., and Berger, I. (2017) MultiBac: from protein complex structures to synthetic viral nanosystems. BMC Biol 15, 99

22. Weissmann, F., Petzold, G., VanderLinden, R., Huis In ’t Veld, P. J., Brown, N. G., Lampert, F., Westermann, S., Stark, H., Schulman, B. A., and Peters, J. M. (2016) biGBac enables rapid gene assembly for the expression of large multisubunit protein complexes. Proc Natl Acad Sci U S A 113, E2564–2569

